# Ensemble AnalySis with Interpretable Genomic Prediction (EasiGP): Computational Tool for Interpreting Ensembles of Genomic Prediction Models

**DOI:** 10.1101/2025.05.25.656055

**Authors:** Shunichiro Tomura, Melanie J. Wilkinson, Owen Powell, Mark Cooper

## Abstract

Ensemble of multiple genomic prediction models have grown in popularity due to consistent prediction performance improvements in crop breeding. However, technical tools that analyse the predictive behaviour at the genome level are lacking. Here, we develop a computational tool called Ensemble AnalySis with Interpretable Genomic Prediction (EasiGP) that uses circos plots to visualise how different genomic prediction models quantify contributions of marker effects to trait phenotypes. As a demonstration of EasiGP, multiple genomic prediction models, spanning conventional statistical and machine learning algorithms, were used to infer the genetic architecture of days to anthesis (DTA) in a maize mapping population. The results indicate that genomic prediction models can capture different views of trait genetic architecture, even when their overall profiles of prediction accuracy are similar. Combinations of diverse views of the genetic architecture for the DTA trait in the TeoNAM study might explain the improved prediction performance achieved by ensembles, aligned with the implication of the Diversity Prediction Theorem. In addition to identifying well-known genomic regions contributing to the genetic architecture of DTA in maize, the ensemble of genomic prediction models highlighted several new genomic regions that have not been previously reported for DTA. Finally, different views of trait genetic architecture were observed across sub-populations, highlighting challenges for between-population genomic prediction. A deeper understanding of genomic prediction models with enhanced interpretability using EasiGP can reveal several critical findings at the genome level from the inferred genetic architecture, providing insights into the improvement of genomic prediction for crop breeding programs.

**Plain Language Summary:** While an ensemble of genomic prediction models has been applied in crop breeding, the prediction mechanism has not been well-investigated due to the lack of a computational tool to interpret the predictive behaviour. It is critical to investigate prediction models at the genome level to understand how each model quantifies genomic marker effects contributing to the trait genetic architecture. Hence, we developed a computational tool, the Ensemble AnalySis with Interpretable Genomic Prediction (EasiGP), to investigate the genome features and predictive behaviours of the ensemble. Here, we demonstrate the utility of EasiGP using a maize breeding dataset. EasiGP visualised the genetic architecture from diverse interpretable genomic prediction models and identified several well-known key maize genes. EasiGP also revealed several potential new genomic regions for further investigation. EasiGP helps us investigate the trait genetic architecture that can be utilised to benefit crop breeding.

**Core ideas:**

- A new computational tool, EasiGP, was created to interpret multiple genomic prediction models at the genomic level
- EasiGP visualises the inferred trait genetic architecture from multiple genomic prediction models with circos plots
- As a case study, EasiGP highlighted several well-known genes regulating the target trait, days to anthesis
- EasiGP can facilitate the discovery of novel genome regions underlying target traits for further investigation

## 1 INTRODUCTION

The popularity of an ensemble of multiple genomic prediction models (Meuwissen et al., 2001) has grown due to the consistent improvement in prediction performance for crop breeding applications (Wallach et al., 2018; McCormick et al., 2021; Heilmann et al., 2023; Kick and Washburn, 2023; Tomura et al., 2025; Washburn et al., 2025). The ensemble approach can potentially be an alternative option to investigations that focus on identifying the “best” individual genomic prediction models that can achieve the highest prediction performance across various prediction scenarios. This possibility aligns with the implication of the Diversity Prediction Theorem (Page, 2011, 2018; Messina et al., 2025) at the genome level, indicating that the prediction error of the ensemble becomes lower than the mean error of the individual prediction models, provided that the predicted values achieved by the individual prediction models are diverse. Hence, we propose that the interpretation of estimated genomic marker effects from multiple prediction models for the analysis of diversity can be a critical concept in the research of genomic prediction.

Interpretability seeks an explanation of the output values from prediction models by estimating the effect of each feature and the interactions among the features (Molnar, 2020). Interpretable prediction models have been leveraged in a wide range of research fields, such as healthcare (Stiglic et al., 2020; Abdullah et al., 2021; Baniecki et al., 2025), chemistry (Oviedo et al., 2022; Tavakoli et al., 2023) and physics (Wetzel et al., 2025). In crop breeding, conventional genomic prediction models, such as standard genomic best linear unbiased prediction (GBLUP; VanRaden, 2008), can reveal the effect of each feature as the additive effect of each genomic marker and the interaction effects as non-additive effects towards target phenotypes, represented as the trait genetic architecture. The application of machine learning models for genomic prediction has also been investigated in crop breeding, considering their complex non-additive predictive algorithms. Genomic prediction algorithms capturing the interaction effects more effectively are expected to improve prediction performance (Hammer et al., 2006; Cooper et al., 2009; Messina et al., 2018; Yadav et al., 2021; Crossa et al., 2025; Tomura et al., 2025). While numerous studies have evaluated the prediction performance of machine learning models (Heslot et al., 2014; Wang et al., 2015; Montesinos-Lopez et al., 2024), the interpretability of genomic marker effects from machine learning algorithms has not been well-investigated in crop breeding due to the complex prediction mechanisms which are referred to as a “black-box” (Danilevicz et al., 2022; Escamilla et al., 2025). The combination of estimated marker effects from conventional and machine learning genomic prediction models as an ensemble needs to be interpretable at the genomic level to improve the characterisation of trait genetic architectures. Hence, the research community in genomic prediction for crop breeding can benefit from the development of a technical tool that can collectively investigate the predictive behaviour of multiple interpretable genomic prediction models at the genome level.

To compare the estimated genomic marker effects of genomic prediction models, circos plots (Krzywinski et al., 2009) can be constructed as a graphical view. Circos plots can be applied to visualise trait genetic architecture by mapping the estimated marker effects to the corresponding genomic regions and highlighting their interactions (Tessele et al., 2025). For example, the circos plot constructed by Wei et al. (2024) visualised the chromosome locations of key genes for 16 traits identified by genome-wide association studies (GWAS) in rice, with links representing epistatic interactions between genomic marker regions. Other research (Saade et al., 2016; Misra et al., 2017; Song et al., 2020) has also constructed circos plots to effectively visualise GWAS results in crop breeding. The circos plot clearly illustrated the distribution of the key genes throughout the genome and their interactions with other genomic regions.

Here, we developed a computational tool called the Ensemble AnalySis with Interpretable Genomic Prediction (EasiGP) to analyse the genetic architectures identified by multiple interpretable genomic prediction models through circos plots. We use the Teosinte Nested Association Mapping (TeoNAM, Chen et al., 2019) experimental results for the days to anthesis (DTA) trait from Tomura et al. (2025) to demonstrate EasiGP and extend the interpretations to the genomic level. EasiGP was evaluated through three objectives: (1) Comparison of the genetic architectures with key genomic marker regions identified by previous studies, assessing the overlap with the known genetic architecture. (2) Comparison between the inferred genetic architectures of the different genomic prediction models to investigate how genomic marker effects were quantified differently. (3) Comparison of the circos plots for each of the five TeoNAM populations to understand the characteristics of the genetic architecture revealed for each population contributing to the TeoNAM. Together, we anticipate that the results obtained from pursuing these objectives will enhance the interpretability of genomic prediction models at the genome level using EasiGP.

## 2 MATERIALS AND METHODS

### 2.1 Overview of EasiGP

EasiGP has two main components, 1) an ensemble of genomic prediction models and 2) circos plots for visualisation of these models at the genetic level (Figure 1). For the genomic prediction function, *N* individual genomic prediction models and the ensemble of the *N* individual genomic prediction models were iteratively developed under *M* independent prediction scenarios. The prediction scenarios were generated by the combination of the total number of sub-populations, the total number of training and test set ratios, the total number of target traits and the total number of random splits of data into the training and test sets. In each prediction scenario, the genotype and phenotype data containing the columns of identification, population, genomic markers and phenotypes with the rows of records were input into the individual genomic prediction models. After the model fitting and genomic prediction using training and test sets, respectively, the individual genomic models returned prediction performance metrics (Pearson correlation and mean square error (MSE)), predicted phenotypes for the training and test sets, genomic marker effects and marker-by-marker interactions. For the ensemble model, the mean predicted phenotype values of the individual genomic prediction models were calculated as the predicted phenotypes of the ensemble model and returned as output of the ensemble model, accompanied by the prediction performance metrics and genomic marker effects.

**Figure 1:**
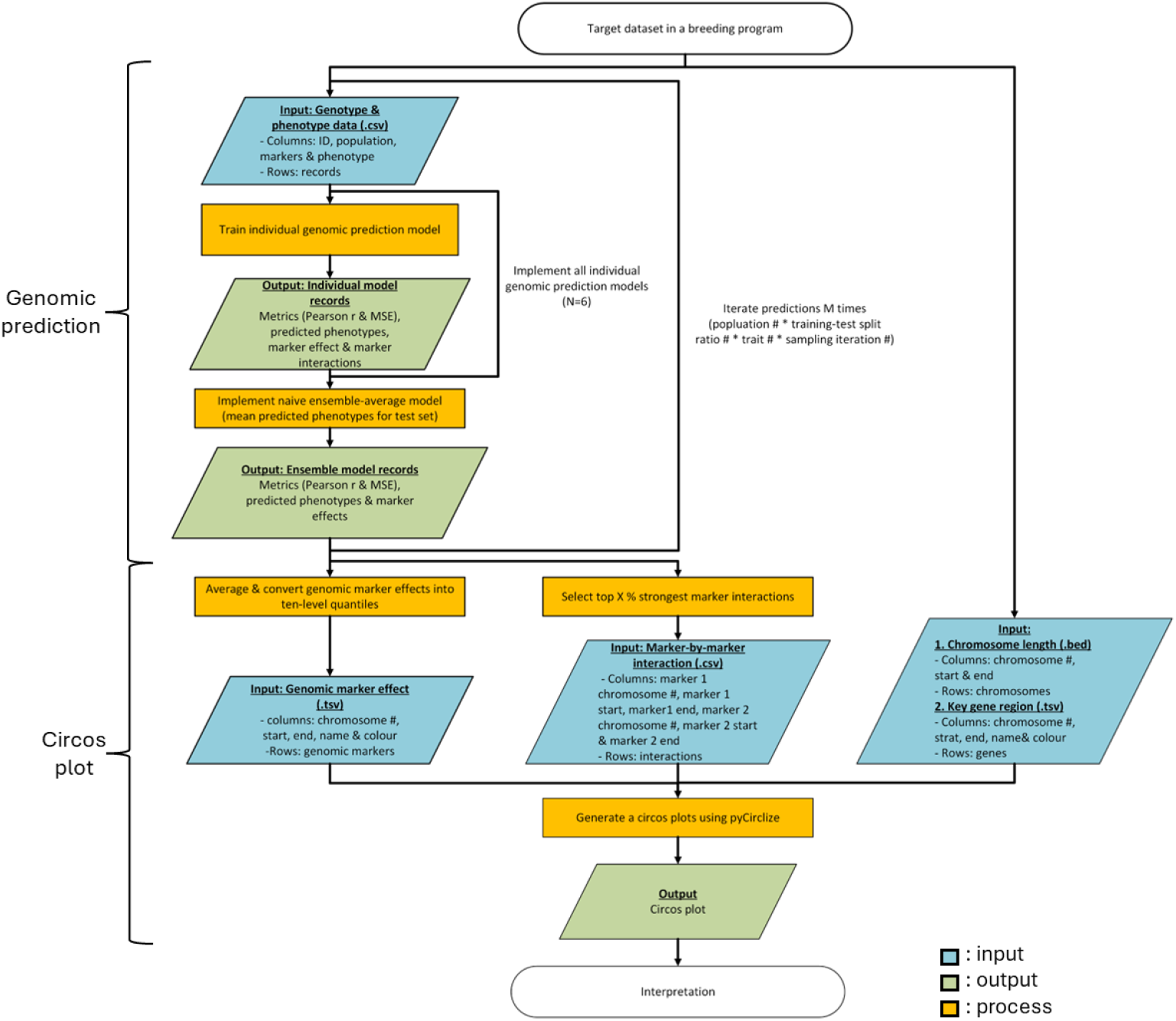
flowchart of EasiGP that consists of two functions; genomic prediction and circos plot development. The blue, green and orange colours represent input, output and process, respectively.

After finishing all the predictive iterations, EasiGP initiates the function for generating a circos plot. The mean genomic marker effects in each prediction model were calculated and converted into the ten-level quantiles. The mean marker-by-marker interaction values were also calculated, and the strongest interactions were selected. Combined with chromosome length and key gene region datasets, a circos plot was generated.

Below, we describe the methods used to estimate the genomic prediction model marker effects and the genomic marker-by-marker interactions and the TeoNAM dataset (Chen et al., 2019) used in this study.

### 2.2 Genomic prediction models

Three classical genomic prediction models (ridge regression best linear unbiased prediction (rrBLUP), BayesB and reproducing kernel Hilbert space (RKHS)) and three machine learning models (random forest (RF), support vector regression (SVR) and graph attention network (GAT)) were applied as the individual genomic prediction models in Tomura et al. (2025). The first five prediction models have been widely used in numerous studies of genomic prediction models. Hence, genome level analysis of the five prediction models may deepen the understanding of the predictive behaviour of commonly used genomic prediction models for crop breeding. For GAT, the prediction mechanism is expected to differ considerably from the other prediction models by incorporating datasets with a graphical representation, potentially generating a different view of the trait genetic architecture, in contrast to the other selected prediction models. EasiGP can be applied to any other possible genomic prediction models to help understand the predictive behaviour at the genome level. The prediction mechanism of each selected genomic prediction model is explained in the subsequent paragraphs.

For the genomic prediction models, the base linear mixed model can be defined as follows (Pérez and de los Campos, 2014):

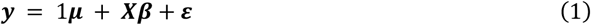

where ***y*** = {*y*_1,_*y*_1,_…,*y*_*n*_} is the set of predicted phenotypes, ***μ***is the intercept, ***X*** is the design matrix of genomic marker values, ***X*** is the allele substitution effect vector of the genomic markers in ***X*** and ***ε***= {*ε*_1,_*ε*_1,_…,*ε*_*n*_} is the vector of residuals. In rrBLUP (Meuwissen et al., 2001), ***β*** is defined as 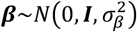 where ***I*** is the identity matrix and 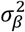 is the variance of genomic marker effects (Endelman, 2011), indicating that ***β*** is assumed to be normally distributed. For BayesB (Meuwissen et al., 2001), on the other hand, some values in ***β*** are diminished to zero with the assumption that not all the genomic markers *J* affect the trait phenotype. For RKHS regression (Gianola et al., 2008), the term ***Xβ*** in Equation (1) is replaced with a term ***μ*** that explicitly captures the non-additive genomic marker effect by projecting the genomic markers into the Hilbert space using a kernel (del los Campos et al., 2009; Pérez and de los Campos, 2014):

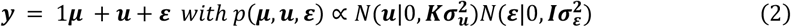

where ***μ*** are the vectors of random effects that are normally distributed with the variance based on the Gaussian kernel matrix 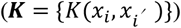. The kernel calculates the mean squared Euclidean distance between genotypes (*x*_*i*_ and 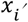):

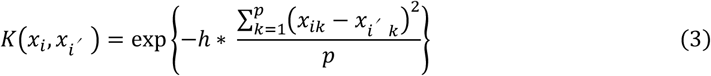

where *h* is the bandwidth parameter that determines the speed of reduction in the covariance function as the distance between the genotype pair increases and *p* is the total number of genomic markers.

For the machine learning models, each aimed to capture both additive and non-additive genomic marker effects through the training process to learn key predictive patterns. RF (Breiman, 2001) is a collection of decision trees that predict target values by tracing the binary decision mechanism using feature values. The final prediction value is calculated by averaging the predicted values from each decision tree. SVR (Drucker et al., 1996) determines the optimal location of a predictive hyperplane by including as many data points as possible within the decision boundary assigned to each side of the hyperplane. The prediction from the optimised hyperplane is expected to minimise the prediction error. GAT (Veličković et al., 2018) is one of the deep learning approaches that leverages graph-structured data to predict target values. Each genomic marker and each phenotype data point is represented as a node, and the genomic marker nodes and phenotype nodes are connected with edges directed from the genomic marker nodes to the phenotype nodes (Figure S1). The GAT application in Tomura et al. (2025) assigned no edges between the genomic marker nodes. This graph structure indicates that each genomic marker affects phenotypes independently and infinitesimally. GAT leverages the attention mechanism that allocates heavier weights to the genomic marker nodes that strongly affect phenotypes, expecting to increase prediction performance.

The naïve ensemble-average model (Tomura et al., 2025) calculated the mean of the predicted phenotypes from each individual genomic prediction model with equal weight. This ensemble model was implemented after each individual genomic prediction model predicted phenotypes.

The genomic prediction models were implemented in R (v1.1.4) and Python (v3.11.10) using software and packages. For rrBLUP, BayesB and RKHS, the software tool BGLR (Pérez and de Los Campos, 2014) was leveraged for the implementation in R. The iteration number and burn-in were set as 12,000 and 2,000, respectively. A Gaussian kernel was used for RKHS. The other parameters were set to default. An increase in the values for the iteration number and burn-in did not change the prediction performance of rrBLUP, BayesB and RKHS. Hence, the provided values were selected considering the computational time. For RF and SVR, Sklearn (v1.2.2) was used to develop the models in Python. The total number of decision trees was 1,000 in RF. All the other hyperparameters such as the maximum depth of each tree, the maximum number of features and the maximum number of samples, were set to default. The radial basis function was selected for the kernel in SVR. The rest of the parameters were set to default. For GAT, PyTorch Geometric (v2.6.1) (Fey et al., 2019) was employed for the implementation in Python. GAT was constructed with one hidden layer containing 20 channels and a dropout of 0. The Exponential Linear Unit function was leveraged as the activation function. GAT was trained iteratively 50 times with the number of batches set as 8. The Adaptive Moment Estimation with Weight Decay (AdamW, Loshchilov and Hutter, 2017) was adopted as the optimiser. The learning rate and weight decay for the optimiser were 0.005 and 0, respectively. The other hyperparameters were set to default.

### 2.3 Genomic marker and marker-by-marker interaction effect inference

Genomic marker effects were inferred from each genomic prediction model to analyse the estimated additive genomic marker effects. For rrBLUP and BayesB, the allele substitution effect ***β*** was used as the genomic marker effect. For RKHS and SVR, the genomic marker effect was estimated using Shapley values (Shapley, 1953) as below (Lundberg and Lee, 2017):

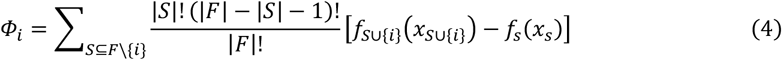

where *Φ*_*i*_ is the Shapley value of the feature *i* (genomic marker), *S* is a subset of the feature collection *F, f*_*S*∪{*i*}_ is the prediction from a model with a feature subset including *i, f*_*s*_ is the prediction from the model without including the feature subset *w*. Shapley values are the cumulative effect of a particular feature, considering all possible combinations based on the conditional probability. In this study, the cumulative effect of each genomic marker on the target phenotype is considered the additive genomic marker effect. Shapley values were calculated for every genotype data point (RIL record) and hence the mean Shapley values for the RILs in the test set were calculated as the final Shapley value. For the implementation, iml (v0.11.3) (Molnar, 2020) was used in R to calculate Shapley values for RKHS. Shapley values were estimated from RKHS due to the inability to directly extract genomic marker effects in contrast to rrBLUP and BayesB. SHAP (v0.42.1) (Lundberg and Lee, 2017) was used for SVR in Python. For RF genomic marker effects, feature importance was leveraged using the impurity-based approach (Ishwaran, 2015). This approach calculates how well each feature (genomic marker) can separate data points containing similar target values into the same nodes in the constituent decision trees. For GAT, Integrated Gradients was leveraged as below (Sundararajan et al., 2017):

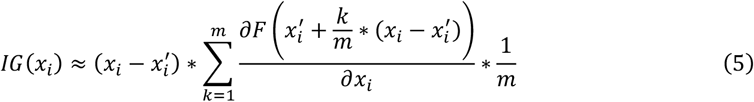

where *x* is a feature (genomic marker), *x*^*′*^ is the baseline value of *x* and *m* is the total number of interpolation steps for the integral. Integrated Gradients calculates the effect of features based on the changes in predicted values between the inclusion and exclusion of the features in the model. Integrated Gradients was calculated per data point (RIL record) and the mean values from the test set were estimated as the final Integrated Gradients. PyTorch Geometric was leveraged for the implementation. The genomic marker effects were inferred in each prediction scenario for every genomic marker. Hence, the mean of the genomic marker effects across all the prediction scenarios was leveraged as the final genomic marker effect.

Using the genomic marker effects estimated from all the individual genomic prediction models, the genomic marker effects for the naïve ensemble-average model were calculated. The genomic marker effects from each individual genomic prediction model were normalised per prediction scenario, and the arithmetic mean of the normalised genomic marker effects across the individual genomic prediction models was used as the genomic marker effects for the naïve ensemble-average model.

For the genomic marker-by-marker interaction effects, pairwise Shapley values were calculated from RF (Equation (3)) using SHAP. Shapley values are calculated by summing all possible feature combinations using conditional probability. To capture the non-additive interactions between genomic markers (features), we combined the effect of a pairwise combination of markers on the target phenotype. The mean genomic marker interaction effects across all prediction scenarios were used as the final genomic marker interaction effects.

### 2.4 Circos plot construction

The circos plots constructed in this study consist of three major sections: (1) key genomic marker region rings representing prior knowledge from previous mapping studies, (2) genomic marker effect rings from genomic prediction models, and (3) genomic marker-by-marker interaction effect links. Each ring consists of segments for the chromosomes of the target species. For the maize TeoNAM (Chen et al., 2019) example in this study, ten segments were developed to represent the total number of chromosomes in maize. The segments were displayed in centimorgans (cM). Circos plots were generated using pyCirclize (v1.7.1) in Python.

#### Prior knowledge

Three ring components were constructed using information from two previous studies. The first ring represented QTL regions identified by Chen et al. (2019). The QTL regions relevant to DTA were inferred and mapped to their corresponding genomic marker regions on the ring. The second and third rings indicated the genomic regions of key genes that operate in the leaf and shoot apical meristem (SAM) tissues to determine the flowering time in maize, respectively (Dong et al., 2012).

#### Genomic Marker effects

Seven ring components were developed to illustrate the genomic marker effects as features of genetic architecture for each genomic prediction model. The inner six rings represented the estimated genomic marker effects from the individual genomic prediction models mapped to the corresponding genome regions. The outermost ring displayed the estimated genomic marker effects for the naïve ensemble-average model (Tomura et al., 2025). The effect sizes of the genomic markers were ranked based on the strength of the corresponding genomic marker effects and split into ten quantiles. The colour intensity indicated marker strength, where darker colours represented larger marker effects. The displayed marker regions on the circos plots were extended by adding 0.2 cM to each side of the actual genomic marker regions for visualisation.

#### Genomic Marker-by-Marker Interactions

The top 0.01% of the most influential genomic marker-by-marker pairwise Shapley scores were extracted from RF as selected interactions and displayed as lines connecting the markers. The strength of the genomic marker interactions was indicated using the width of the corresponding links; thicker links were assigned to stronger genomic marker interactions. The genomic marker interactions were classified into two categories: genomic marker interactions within the same chromosome and those between different chromosomes.

### 2.5 Dataset

The TeoNAM dataset (Chen et al., 2019) contains the genotype and phenotype records of recombinant inbred lines (RILs) from the five hybrid sub-populations between the W22 maize (*Zea mays*) line and five teosinte types (*Z. mays ssp. parviglumis* for TIL01, TIL03, TIL11 and TIL14 and *Z*.*mays ssp. Mexicana* for TIL25). In each RIL sub-population, the F1 generation was developed by crossing W22 and the teosinte type, backcrossed with W22 in the subsequent generation. Selfing was then conducted for the next four generations. Each RIL sub-population was evaluated twice at the University of Wisconsin West Madison Agricultural Research Station in a randomised complete block. The total number of genomic markers and records from each of the five RIL sub-populations is summarised in Table S1.

The diverse crosses with Teosinte motivated the selection of the TeoNAM dataset for this study to demonstrate the application of EasiGP. The crosses with an ancestral species of maize can diversify the fixed gene networks for traits in the domesticated maize lines. The increased genetic diversity within the TeoNAM crosses may help the genomic prediction models identify key genomic marker regions and pathways.

The days to anthesis (DTA) trait was selected in this study primarily due to previous studies revealing key genomic regions contributing to DTA (Buckler et al. 2009; Dong et al., 2012; Chen et al., 2019; Wisser et al., 2019). This prior knowledge can be used to assess whether the genomic prediction models used in this study can correctly detect known causative loci expected to segregate in the TeoNAM populations. A deeper understanding of the genetic architecture of DTA using the interpretable genomic prediction models can help improve the selection of genomic markers for predicting DTA, ultimately benefiting prediction models used for crop yield improvement.

Each RIL contains two phenotype scores for DTA, since each population was evaluated twice, and hence the TeoNAM dataset consists of two sub-datasets in each population record. Two sub-datasets were concatenated into one within each population record, with a factor explicitly representing different evaluations with the value of 0 or 1.

Three data preprocessing methods were applied to the TeoNAM dataset prior to training the genomic prediction models in Tomura et al. (2025): missing allele imputation for genomic markers, missing trait phenotype RIL removal and the reduction of genomic markers based on linkage disequilibrium (LD) relationships. The missing alleles were imputed using the most frequent allele of the population at that site after excluding genomic markers containing more than 10% of missing marker calls. All the RILs with a missing phenotype value were excluded from the dataset. LD relationships between markers were leveraged to mitigate the effect of the curse of dimensionality (Bellman, 1957; Ramstein et al., 2019) by reducing the total number of genomic markers. LD filtering was performed using PLINK (v1.9) (Chang et al., 2015) with a window size of 30,000, a step size of 5 and a squared correlation (*r*^*2*^) of greater than 0.8.

In this study, each sub-population was randomly split into a training and test set 500 times with three different training and test set ratios (0.8-0.2, 0.65-0.25 and 0.5-0.5). Hence, the genomic marker and interaction effects were calculated for 7,500 different prediction scenarios (5 sub-populations * 3 ratios * 500 random split sampling iterations) for each genomic prediction model. For the circos plot development, the mean genomic marker and interaction effects across all the 7,500 prediction scenarios were leveraged to infer the genetic architecture by constructing the genomic marker effect rings and genomic marker-by-marker interaction effect links in the circos plots.

For instance, the genomic prediction function in EasiGP may initiate from targeting the prediction of DTA in population W22TIL01 with the training-test split ratio of 0.8-0.2. The preprocessed W22TIL01 sub-population data, containing 444 records and 274 genomic markers, is randomly split into training (355 records) and test sets (89 records) as the initial procedure of the phase. After training the genomic prediction models, the effect size of all 274 genomic markers and marker-by-marker interactions is estimated from all the prediction models using the respective genomic marker effect estimation methods described in the previous subsections. Such an estimation process iterates 500 times with different training-test set ratios in each prediction scenario. The mean effect size of each genomic marker and marker-by-marker interaction is calculated in the circos plot development function to represent overall genomic marker effects across different scenarios, subsequently linked to corresponding genomic marker regions to map the genomic marker effects as a circos plot.

## 3 RESULTS

### 3.1 Genomic prediction models identified several key genomic marker regions

The genomic prediction models identified several genomic marker regions that collocate with previously discovered influential genes for DTA in maize (Figure 2; Table 1). One of the strongest overlaps was observed in genomic marker regions containing genes involved in the photoperiod pathways, specifically ZmCCT9, ZmCCT10 and ZCN8. Almost all the genomic prediction models highlighted the genomic marker regions corresponding to these three genes. The word “highlight” here is defined as the genomic marker regions that were continuously in the top 10% of genomic marker effects across multiple prediction models. ZCN8 is a key gene for determining flowering time in maize, controlling the transition from the vegetative to reproductive stage by functioning in both the leaf and SAM (Meng et al., 2011; Dong et al., 2012; Guo et al., 2018; Aiyesa et al., 2025). ZmCCT9 promotes flowering time when daylight hours become longer at high-latitude regions by regulating ZCN8 (Huang et al., 2018a). ZmCCT10 is another critical gene in the photoperiod pathway, which delays flowering under long days through interactions between ZCN8 and ZmCCT9 (Dong et al., 2012; Hung et al., 2012; Wisser et al., 2019). Such close interactions between the genes in the photoperiod pathway were captured in the circos plot by showing heavy marker-by-marker interactions between the regions of those genes, especially between chromosome 8 (ZCN8) and 10 (ZmCCT10) (Figure 2).

**Table 1:** A list of key genomic marker regions highlighted as strong features by at least two genomic prediction models. The gene names and functions were based on Dong et al. (2012) and Chen et al. (2019). Genomic prediction model indicates the models that continuously allocated the top 10% of genomic marker effects to the corresponding key genomic marker regions.

| Chromosome | Region (cM) | Gene name | Function | Genomic prediction model |
| --- | --- | --- | --- | --- |
| 1 | 33 - 36 | PhyB1 | Light transduction | rrBLUP, BayesB, RKHS, GAT and Ensemble |
| 1 | 80 - 85 | - | - | rrBLUP, BayesB and RF |
| 2 | 10 - 15 | ZFL2 | Integrator | rrBLUP, BayesB, and GAT |
| 2 | 133 - 134 | FCA | Autonomous | RKHS, GAT and Ensemble |
| 3 | 5 - 10 | miR156, GIGZ1B | Aging, Clock | rrBLUP, BayesB, RKHS, SVR, GAT and Ensemble |
| 3 | 85 - 90 | ZmMADS69 | Adaptation | All |
| 4 | 55 - 60 | ZmLHY2 | Clock | All |
| 6 | 75 - 76 | ZAG1 | Flower development | BayesB, RKHS, RF, SVR, GAT and Ensemble |
| 7 | 68 - 70 | - | Early flowering | RKHS, GAT and Ensemble |
| 7 | 140 - 145 | dlf1, ZmPRR37 | Integrator, Clock | All |
| 8 | 65 - 70 | ZCN8 | Photoperiod | rrBLUP, BayesB, RF, SVR, GAT and Ensemble |
| 9 | 70 - 75 | ZmCCT9 | Photoperiod | rrBLUP, BayesB, RF, SVR, GAT and Ensemble |
| 10 | 40 - 55 | ZmCCT10 | Photoperiod | rrBLUP, BayesB, RF, SVR, GAT and Ensemble |
| 10 | 110 -115 | ZFL1 | Integrator | RKHS, RF and Ensemble |

**Figure 2:**
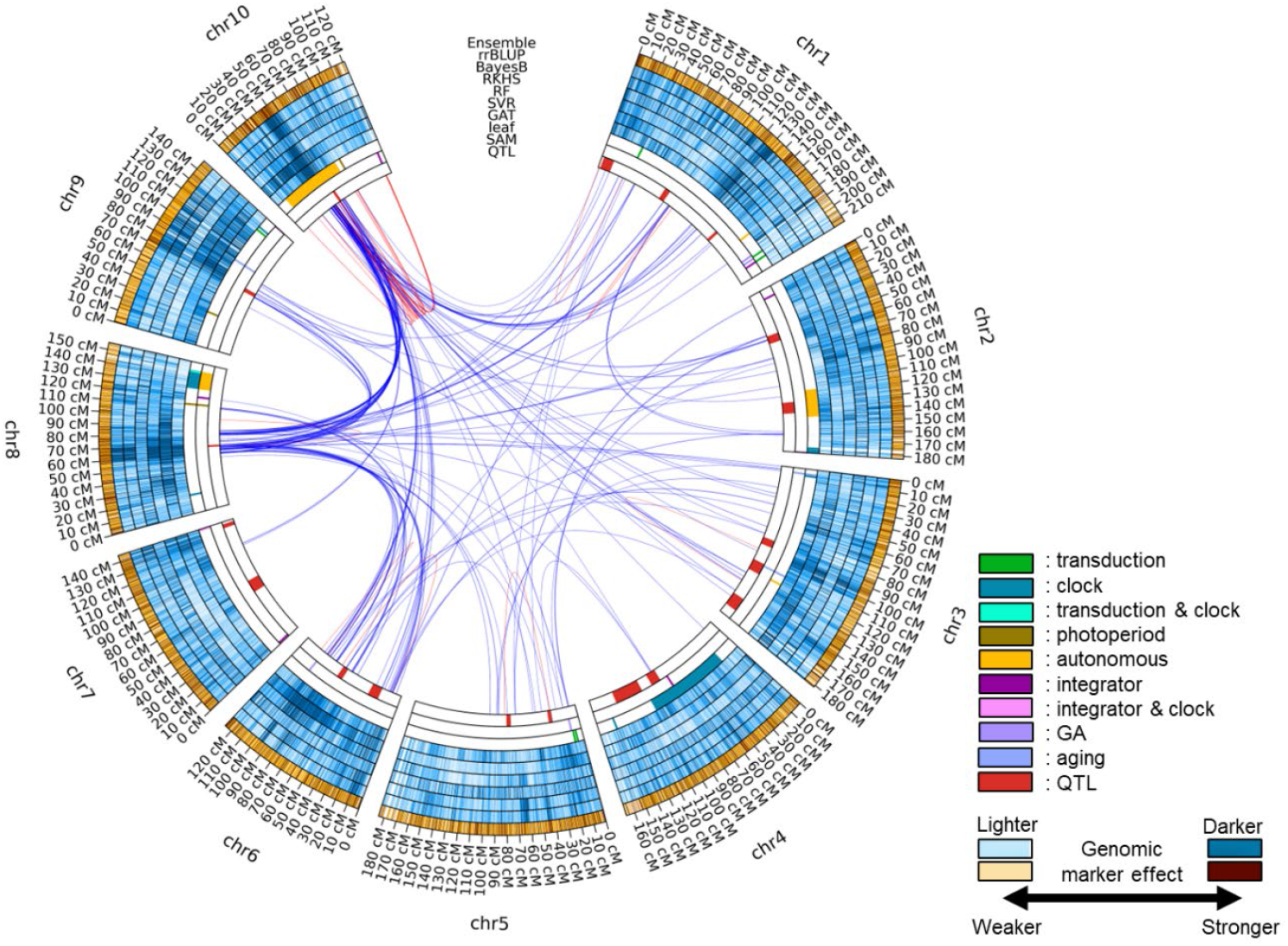
Circos plot for the trait days to anthesis (DTA) in the TeoNAM dataset across the five populations. The innermost ring (QTL) has the QTL genomic marker regions for DTA identified in Chen et al. (2019). The second and third innermost rings (SAM and leaf) represent the key genomic marker regions that affect the shoot apical meristem (SAM) and leaf, respectively, from Dong et al. (2012). These key genomic marker regions affect light transduction (transduction), circadian clock (clock), photoperiod, autonomous, integrator, gibberellin (GA) and aging pathways. The subsequent blue rings (fourth to ninth) represent the genomic marker effects of each genomic marker region inferred from the GAT, SVR, RF, RKHS, BayesB and rrBLUP genomic prediction models, respectively. The outermost orange ring is the genomic marker effect estimated for the naïve ensemble-average model. The intensity of blue and orange colours shows the strength of the genomic marker effect classified into ten quantiles. Darker colours are allocated to the genomic marker regions with higher genomic marker effect levels. The red and blue lines between genomic marker regions are the genomic marker-by-marker interaction effects estimated by pairwise Shapley values from RF (top 0.01%; red = within chromosome and blue = between chromosomes).

Several colocation overlaps were also identified at the genomic marker regions regulating the circadian clock (ZmLHY2, ZmPRR37 and GIGZ1B). ZmLHY2 is a transcription factor that regulates the morning loop of the circadian clock (Hayes et al., 2010). ZmPRR37 also regulates the circadian clock in maize by signalling diurnal outputs, which is a homolog to PRR genes in rice (Hayes et al., 2010). GIGZ1B is expressed at its peak level during the daylight hours, controlling the circadian clock, regardless of the length of daylight (Miller et al., 2008). These colocation overlaps of regions identified by the genomic prediction models with regions containing known genes involved in controlling maize DTA indicate that the genomic prediction models captured several key features of genomic marker information relating to the circadian clock.

Additionally, the genomic prediction models identified several key genomic marker regions in chromosome 2, 7 and 10 that contain genes that function as integrators of the different pathways (ZFL1, ZFL2 and dlf1). These integrators respond to the signals from the multiple pathways to determine DTA as an output of the regulatory network. Hence, integrators may contain multiple different functions. ZFL1 and ZFL2 are involved in the transition from the vegetative stage to the reproductive stage by upregulating the ABC floral organ identity network of genes (Bomblies et al., 2003). In addition, ZFL2 is known to control tassel branching and the traits relating to inflorescence structure in maize (Bomblies and Doebley, 2006). dfl1 encodes a protein with a basic leucine zipper domain (Muszynski et al., 2006), expressed in the leaves and regulating the timing of the transition to the flowering stage by sending signals from leaves to the SAM.

Other key genomic marker regions identified by the genomic prediction models, such as PhyB1, FCA, miR156, ZmMADS69 and ZAG1, influence the flowering time in maize by affecting light transduction, aging, adaptation and flower development pathways. Collectively, those results indicate that the genomic prediction models captured several key genomic marker regions and interactions known to be involved in the genetic architecture of DTA in maize.

The genomic prediction models also highlighted several other genomic marker regions that have not been well-studied previously (Figure 2). For instance, while the genomic marker region in chromosome 1 (from 125 cM to 135 cM) was highlighted by all the individual genomic prediction models besides RKHS, no nearby key genomic marker regions were identified in the previous studies. Similarly, another genomic marker region in chromosome 1 (from 185 cM to 190 cM) was highlighted in the inferred genetic architecture, except for RKHS and SVR, but no key genomic marker region overlapped. Such discordant genomic marker regions were observed in several other genomic marker regions (from 85 cM to 90 cM in chromosome 2 and 45 cM and 65 cM in chromosome 5). Those highlighted genomic marker regions indicate that the genomic prediction models heavily weighted the predictive information from several genomic marker regions not identified in the previous studies.

### 3.2 Each genomic prediction model captured different genetic architectures

While the genomic prediction models identified several key genomic marker regions, their relative magnitudes differed among the genomic prediction models. The different relative values are indicated by the unique patterns of colour intensity for the deciles within each ring (Figure 2). For example, while the genomic marker region between 30 cM and 60 cM in chromosome 10 showed a similar pattern in the colour intensity for rrBLUP, BayesB, RF and GAT, the genetic architecture of RKHS and SVR in this region showed a unique pattern. RKHS did not specifically highlight the region as highly influential, while SVR emphasised the genomic marker effects on the first half of chromosome 10 more than the other genomic prediction models. Such diversity was also observed in the other highlighted genomic marker regions that overlapped with key genomic marker regions identified by the previous studies (Table 1). While several key genomic marker regions were identified across all the genomic prediction models as contributing strong marker effects, such regions were mainly highlighted by a fraction of the genomic prediction models. The lack of a clear consensus in the highlighted genomic marker regions for all six genomic prediction models considered also showed that each genomic prediction model quantified genomic marker effects differently. Such uniqueness led to different views of the genetic architecture of DTA in the TeoNAM dataset for the different genomic prediction models.

### 3.3 Inferred genetic architectures differed among the TeoNAM sub-populations

The patterns of the quantified genomic marker effects in each genomic marker region were distinctive in each of the five sub-populations composing the TeoNAM dataset (Figure 3). For instance, while the genomic marker region containing ZmCCT10 was highlighted in chromosome 10 across the sub-populations, the range of the highlighted region varied among sub-populations. For W22TIL01 and W22TIL25, the highlighted region was between 40 cM and 50 cM (Figure 3a, 3e), whereas the region was extended in W22TIL03 (between 45 cM and 55 cM) and W22TIL14 (between 30 cM and 55 cM) (Figure 3b, 3d). The region for W22TIL11 was between 40cM and 55cM (Figure 3c) with a lower frequency of highlights compared to the others. Similarly, the genomic marker region between 85 cM and 95 cM in chromosome 1 was mainly allocated to the top 10% of the genomic marker effects by all the genomic prediction models in W22TIL03 (Figure 3b). However, the genomic prediction models in the other four sub-populations did not frequently allocate such high genomic marker effects to those regions of the genome. Instead, the genomic prediction models more frequently assigned higher genomic marker effects to other genomic marker regions. Further, while all the genomic prediction models frequently assigned the top 10% of the genomic marker effects to the genomic marker region between 60 cM and 80 cM in chromosome 9 of W22TIL03 (Figure 3b), the genomic prediction models did not clearly highlight the same genomic marker region in the four other sub-populations. A lack of consensus in the quantification patterns of the genomic marker effect among the five TeoNAM sub-populations was observed for other genomic marker regions across the chromosomes. These diverse patterns indicate that the genomic prediction models recognise different genomic marker regions as influential in each of the five sub-populations comprising the TeoNAM dataset.

**Figure 3:**
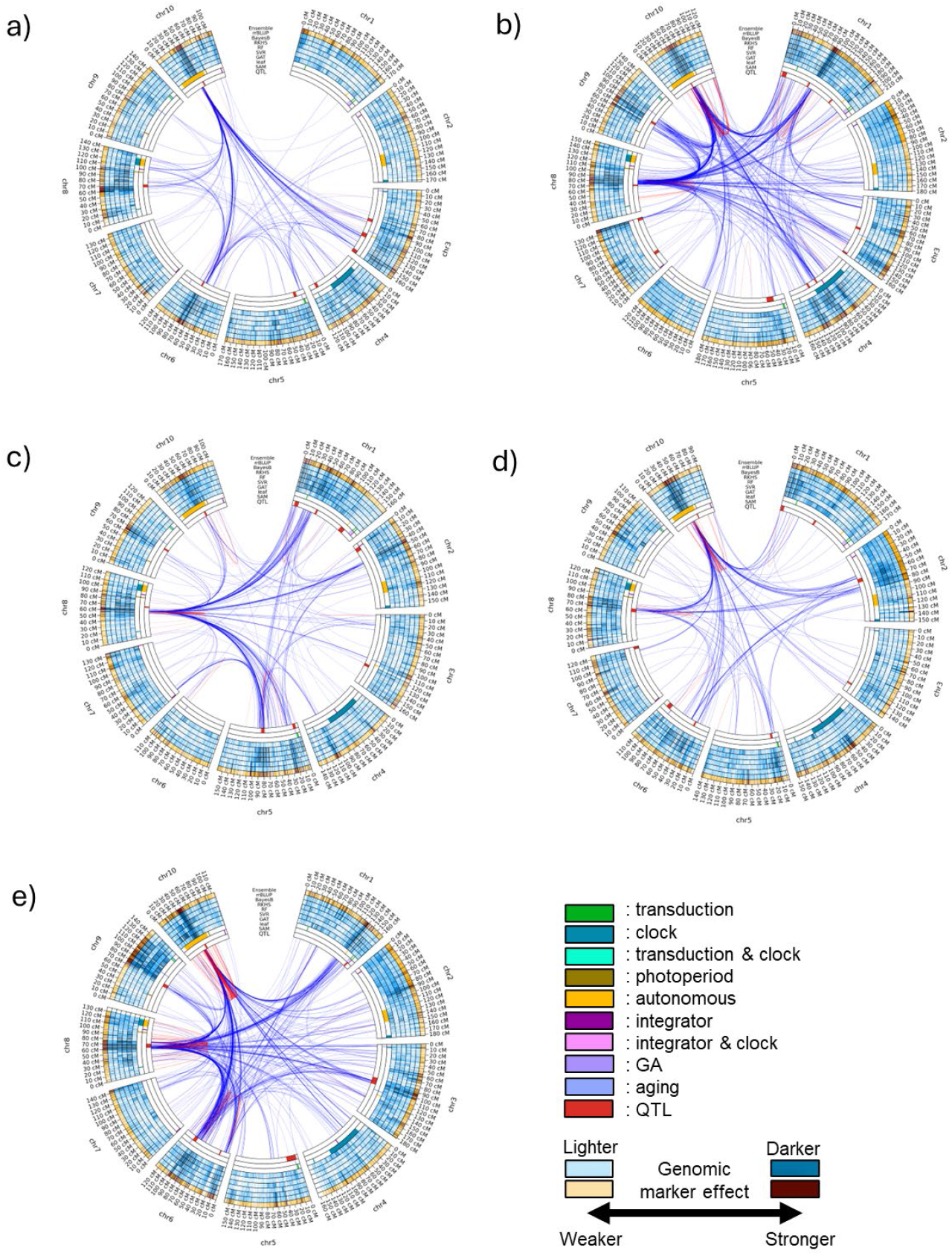
Circos plots for days to anthesis (DTA) in the TeoNAM dataset for the five sub-populations: a) W22TIL01, b) W22TIL03, c) W22TIL11, d) W22TIL14 and e) W22TIL25. The innermost ring (QTL) has the QTL genomic marker regions identified in Chen et al. (2019). The second and third innermost rings (SAM and leaf) represent the key genomic marker regions that affect the shoot apical meristem (SAM) and leaf, respectively, from Dong et al. (2012). These key genomic marker regions affect light transduction (transduction), circadian clock (clock), photoperiod, autonomous, integrator, gibberellin (GA) and aging pathways. The subsequent blue rings (fourth to ninth) represent the genomic marker effects of each genomic marker region inferred from GAT, SVR, RF, RKHS, BayesB and rrBLUP, respectively. The outermost orange ring is the genomic marker effect estimated from the naïve ensemble-average model. The intensity of blue and orange colours shows the strength of the genomic marker effect classified into ten levels based on the quantiles. Darker colours are allocated to the genomic marker regions with higher genomic marker effect levels. The red and blue lines between genomic marker regions are the genomic marker interaction effect estimated by pairwise Shapley values from RF (top 0.01%; red = within chromosome and blue = between chromosomes).

Such diversity in the marker effect combinations among the six genomic prediction models for each sub-population was also observed in the inferred genomic marker-by-marker interaction effects. While the strong interactions between chromosomes 8 and 10 were conserved across the sub-populations, many of the detected marker-by-marker interaction regions differed among the five sub-populations. The region for chromosome 8 was located at 70 cM in W22TIL01, W22TIL14 and W22TIL25 (Figure 3a, 3d, 3e), whereas the region was located at 80 cM in W22TIL03 and 60 cM in W22TIL11 (Figure 3b, 3c). For chromosome 10, the region was located between 40 cM and 50 cM in W22TIL01 and W22TIL25 (Figure 3a, 3e), between 50 cM and 60 cM in W22TIL03 (Figure 3b), between 45 cM and 55 cM in W22TIL11 (Figure 3c) and between 30 cM and 45 cM in W22TIL14 (Figure 3d). Additionally, this region in chromosome 8 strongly interacted with other chromosomes in unique patterns identified across the five sub-populations. While all the sub-populations showed interactions with chromosome 5, this was stronger than for the other sub-populations in W22TIL11 (Figure 3c). Three sub-populations (W22TIL03, W22TIL11 and W22TIL25) showed heavy marker-by-marker interactions associated with chromosome 1 and chromosome 3 (Figure 3b, 3c, 3e), whereas there were strong interactions associated with chromosome 2 for W22TIL11 and W22TIL14 (Figure 3c, 3d). Such distinctive marker-by-marker interaction patterns were observed throughout the chromosomes. Those results indicate that the inferred views of trait genetic architecture were diverse across the five sub-populations.

## 4 DISCUSSION

### 4.1 EasiGP helps investigate genomic prediction models at the genome level

The circos plots generated by EasiGP provided a genome level view of how the six investigated genomic prediction models estimated diverse genomic marker effects (Figure 2). This diversity is attributed to algorithmic differences among the prediction models. The algorithms of rrBLUP and BayesB primarily targeted the detection of the additive genomic marker effects (Meher et al., 2022), while RKHS also included a kernel to explicitly identify several non-additive genomic marker-by-marker interaction effects (Gianola et al., 2008). The machine learning models included components in their prediction algorithms to identify more complex non-additive prediction patterns underlying the inclusion of the genetic markers (Chen et al., 2023). The comparison of the estimated feature importance from logistic regression (additive prediction model) and RF (non-additive prediction model) indicated that several identified key features varied between the prediction models for the classification tasks in the cancer study (Saarela and Jauhiainen, 2021). A linear regression model was also compared with RF in regression tasks, showing a similar conclusion that the level of importance in each feature identified by the linear regression was not necessarily aligned with that of RF (Grömping, 2009). The estimated patterns in the feature importance were also investigated within three machine learning models (multilayer perceptron (MLP), naïve Bayes and Bagging), with the observation that each prediction model weighted feature importance distinctively (Štrumbelj and Kononenko, 2014). Hence, the different prediction strategies in each prediction algorithm quantified the genomic marker effects uniquely throughout the chromosomes.

Such differences in genome profiles of the captured genomic marker effects were represented as diverse views of the genetic architecture of the trait DTA in our study using the TeoNAM dataset (Figure 2, Table 1). The circos plot enabled the visualisation of diverse patterns in the colour intensity across the chromosomes that were not identical to any other views. The combination of such diverse views in the ensemble approach provides a novel view of marker weights, taking into consideration the consistent performance improvement of the ensemble approach observed in Tomura et al. (2025). The weaknesses of individual prediction algorithms can potentially be offset by aggregating the diversely quantified genomic marker effects from each model (Kick and Washburn, 2023; Washburn et al., 2025). The interpretation of the visualised genetic architecture for the ensemble is supported by the Diversity Prediction Theorem (Page, 2018), implying that diversified predicted phenotypes through differently quantified genomic marker effects contributed to the reduction of the ensemble error for the naïve ensemble-average model compared to the mean model error. This implication is also aligned with the explanation of the No Free Lunch Theorem (Wolpert and Macready, 1997), stating that no individual prediction model can consistently reach the highest prediction performance across the prediction scenarios. The investigation of the genomic prediction models at the genomic level deepens our understanding of the possible reasons for the prediction performance improvement by the ensemble. The circos plot visualises how an ensemble can leverage the diversity of prediction algorithms to model the high dimensionality of the genotype-to-phenotype state space for complex traits.

The application of both interpretable prediction models and circos plots has been reported in several studies, primarily in studies of the human genome, using the self-attention mechanism and statistical analysis (Lee et al., 2023; Peng et al., 2023; Prakash and Banerjee, 2023). However, this approach has not been widely applied to the genome-level study of genomic prediction models for crop breeding applications. Hence, we have developed EasiGP to enable the construction of circos plots that visualise genomic marker effects from interpretable genomic prediction models and their ensemble.

### 4.2 EasiGP may reveal potential new key genomic marker regions

The circos plot view of the six genomic prediction models and the ensemble highlighted several genomic marker regions that were not previously identified as key genomic regions (Figure 2). Those regions did not overlap with key gene regions reported from previous studies and could potentially be leveraged as new genomic marker regions that can contribute to improving target traits. There are multiple possible reasons why these new regions were identified by the genomic prediction models rather than by traditional QTL mapping. Those reasons can be attributed to the fundamental distinction that traditional QTL analysis leverages different approaches for QTL detection than the interpretable genomic prediction (Holland, 2004; Alonso et al., 2006). This distinction is largely driven by QTL analysis focusing on testing individual loci for significance, whereas interpretable genomic prediction models the entire genome simultaneously.

The use of traditional and new approaches can contribute to identifying more comprehensive sets of key genomic markers for target traits. The combination of both QTL analysis and interpretable prediction models (machine learning) has started being evaluated in crop breeding programs targeting soybeans (Gill et al., 2022; Yoosefzadeh et al., 2022). Since the use of interpretable genomic prediction for identifying key genomic marker regions has been a relatively recent approach in crop breeding, it is vital to verify the reliability through empirical studies and simulations. Nevertheless, there is potential for the collective analysis of key genomic marker regions to enhance the understanding of trait genetic architecture that can support prediction applications for breeding.

### 4.3 Distinctive genetic architecture in each population may hinder accurate between-population prediction

The genomic prediction models identified different key patterns in the genomic marker and marker-by-marker interaction effects across the five sub-populations comprising the TeoNAM dataset (Figure 3). While similar genomic marker regions were highlighted with heavy interactions for some regions, the majority of the genetic architectures identified by the genomic prediction models showed unique patterns by quantifying different genomic marker regions with high genomic marker and marker-by-marker interaction effects. The diversity in the genetic architecture at the sub-population level might have been attributed to the structure of the TeoNAM populations. The development of NAM populations was proposed to improve resolution through diversifying the allele combinations within the mapping study while maintaining common linkage information for QTL analysis, achieved by leveraging a common parent and numerous donor parents (Gage et al., 2020). Through this population design, each population is developed to be genetically diverse from the others. Consequently, the allele combination became unique with different allele frequency patterns in each population (Gireesh et al., 2021). In the TeoNAM dataset, five sub-populations were developed from one of the two teosinte types crossed with the W22 maize line used as the common parent, followed by backcrossing and a series of selfing generations (Chen et al., 2019). Thus, the five sub-populations were characterised by distinctive allele combination patterns. Consequently, the genetic architecture of each sub-population showed a distinctive structure.

The differences in the genetic architecture for each sub-population may hinder the between-sub-population prediction with high accuracy. Each genomic prediction model learns key prediction patterns from the genetic architecture of the training set. However, the learnt prediction patterns may not be aligned with the key prediction patterns in the test set from another sub-population due to differences in the standing genetic variation. For example, in wheat, the prediction performance of the between-population prediction for 19 traits was considerably lower than the within-population prediction (Huang et al., 2018b). This poor prediction performance was attributed to a lack of lines closely related to the test set. The performance of the between-population prediction for sorghum depended on the combination of the populations for the training and test sets (Sapkota et al., 2020). These experimental results from previous studies indicate that the differences in the genetic architecture between the training and test sets considerably affect the performance of genomic prediction models for the between-population scenarios. Higher prediction performance was achieved when the population structure between the training and test sets was similar. De Roos et al. (2009) concluded that the performance of between-population prediction increases when samples from all populations are included in the training set. The inclusion of samples from different populations allows genomic prediction models to capture multiple views of the trait genetic architecture.

## 5 CONCLUSIONS

EasiGP supports visualisation of the inferred genetic architectures from the six interpretable individual genomic prediction models reported in Tomura et al. (2025) through an application of circos plots. The generated circos plots revealed important diversity in the trait genetic architectures identified for the different genomic prediction models by quantifying genomic marker effects differently while still capturing several key genomic marker regions. The aggregation of such diverse views can potentially enhance the prediction performance using the ensemble, as indicated by the Diversity Prediction Theorem. The diverse views of the trait genetic architecture across TeoNAM sub-populations indicate a source of difficulty in achieving high accuracy for between-population predictions. There is also potential for the interpretable genomic prediction model approach to discover novel key genomic marker regions that have not been identified previously. Through application to appropriate training sets, EasiGP can deepen the understanding of each interpretable genomic prediction model at the genome level and thus benefit crop breeding programs by deepening our understanding of trait genetic architecture and the prediction performance improvement using the ensembles of genomic prediction models.

## Abbreviations

AdamW: adaptive moment estimation with weight decay
DTA: days to anthesis
EasiGP: Ensemble AnalySis with Interpretable Genomic Prediction
F1: filial 1
GAT: graph attention network
GBLUP: genomic best linear unbiased prediction
GWAS: genome-wide association study
LD: linkage disequilibrium
MLP: multi-layer perceptron
NAM: nested association mapping
QTL: quantitative trait locus
RF: random forest
RIL: recombinant inbred line
RKHS: reproducing kernel Hilbert space
SAM: shoot apical meristem
SNP: single nucleotide polymorphism
SVR: support vector regression
TeoNAM: teosinte nested association mapping

## DATA AND CODE AVAILABILITY

EasiGP is available at https://github.com/ShunichiroT/EasiGP. The original TeoNAM dataset (Chen et al., 2019), leveraged to infer the genomic marker and interaction effects, is available at https://datac-ommons.cyverse.org/brow-se/iplant/home/shared/panzea/genotypes/GB-S/TeosinteNAM for the genotypes and https://gsajournals.figshare.com/articles/dataset/Supplemental_Material_for_Chen_et_al_2-019/9250682 for the phenotypes.

## AUTHOR CONTRIBUTIONS

Shunichiro Tomura: Formal analysis, Investigation, Methodology, Software, Validation, Visualisation, Writing – original draft and Writing – review & editing; Melanie J. Wilkinson: Validation, Supervision and Writing – review & editing; Owen Powell: Investigation, Supervision and Writing – review & editing; Mark Cooper: Conceptualization, Investigation, Methodology, Funding acquisition, Project administration, Resources, Supervision, Validation, Writing – review & editing

## ACKNOWLEDGEMENTS

We thank the National Computing Infrastructure (NCI) and Research Computing Centre (RCC) at the University of Queensland for providing access to use High Performance Computing (HPC) machines. We also thank Mr. Greg McLean for constructive feedback to improve the implementation of EasiGP on GitHub.

## Supplementary materials

**Table S1:**
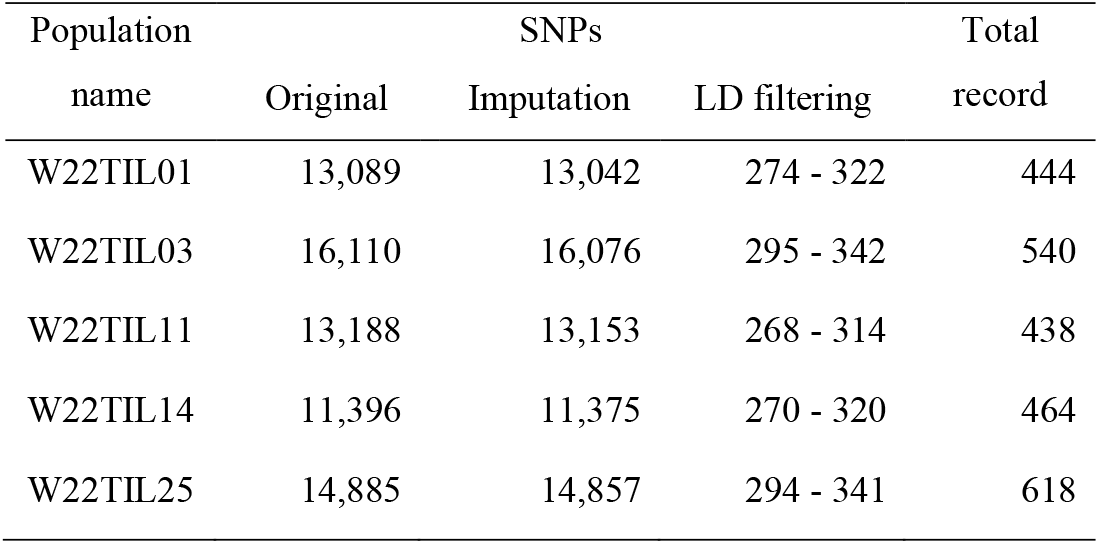
The total number of genomic markers (SNPs) and the total number of records obtained from recombinant inbred lines (RILs) scored twice in each population, created from Tomura et al. (2025). “Imputation” indicates the total number of SNPs after excluding SNPs containing more than 10% of missing marker calls during the missing allele imputation process with the most frequent alleles. Subsequently, remaining SNPs were filtered through linkage disequilibrium (LD) with a threshold of 0.8. The total number of final SNPs after the “LD filtering” depends on the combination of RILs included in the training set in each prediction scenario.

**Figure S1:**
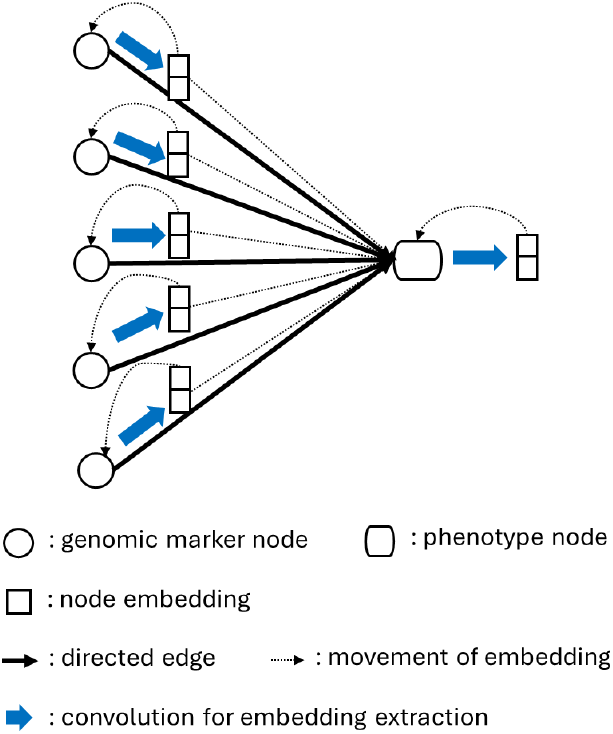
Simplified illustrative view of a graph attention network (GAT) in this study. Each genomic marker node was connected to the phenotype node with an edge directed from the marker nodes to the phenotype node. Key predictive information was extracted from each marker node as an embedding in a vector format through convolution, subsequently gathered with weights as attentions to calculate the embedding of the phenotype node. The embedding of each node was reassigned to the corresponding node as the initial node information for the next embedding extraction (hidden) layer. The embedding size was determined by the number of channels in each hidden layer (2 in this example). At the last (outer) layer, the size of the embedding was set as 1. The embedding of the phenotype node at the last layer is returned from GAT as a predicted phenotype.

